# In-cell structural insight into the stability of sperm microtubule doublet

**DOI:** 10.1101/2023.05.30.542832

**Authors:** Linhua Tai, Guoliang Yin, Xiaojun Huang, Yun Zhu, Fei Sun

## Abstract

The propulsion for mammalian sperm swimming is generated by flagella beating. Microtubule doublets (DMTs) along with microtubule inner proteins (MIPs) are essential structural blocks of flagella. However, the intricate molecular architecture of sperm DMT remains elusive. Here, by *in situ* cryo-electron tomography, we solved the in-cell structure of mouse sperm DMT up to 3.6 Å resolution and built its model with 36 kinds of MIPs in 48 nm periodicity. We identified multiple copies of Tektin5 that reinforce Tektin bundle, and multiple MIPs with different periodicities that anchor the Tektin bundle to tubulin wall. This architecture contributes to a superior stability of A-tubule than B-tubule of DMT, which was revealed by structural comparison of DMTs from the intact and deformed axonemes. Our work provides an overall molecular picture of sperm DMT that is essential to understand the molecular mechanism of sperm motility as well as the related ciliopathies.

## Main

Sperm is a type of cell with specialized function and morphology. It carries the male partner’s genetic material and proteins to find and fuse with the female oocyte at fertilization ^1^. Mature mammalian sperm consists of two main parts: a head with nucleus and acrosome, and a long tail for movement. The tail of sperm is also known as motile cilia or flagella ^2^, and its core component is the axoneme, a microtubule-based super-large molecular machinery ^3^. Inside the sperm axoneme, a pair of microtubule singlets termed as central pair complex (CPC) are surrounded by 9 microtubule doublets (DMTs), forming a ‘9 + 2’ architecture ^4,5^. There are hundreds of different proteins decorated on the microtubule-based structures to form a variety of functional components, such as microtubule inner proteins (MIPs), inner and outer dynein arms (IDA and ODA), radial spokes (RS), and nexin-dynein regulatory complex (N-DRC) ^2^. The dynein motors are responsible for driving sperm movement and fine-tuning motility, while the MIPs provides the major stabilizing force for the DMT architecture ^6^.

In the early 21^st^ century, sperm axoneme was discovered to have a periodicity of multiples of 8 nm with MIPs attached to the inner surface of DMTs by using the approach of cryo-electron tomography (cryo-ET) and sub-tomogram averaging (STA) ^7-11^. Subsequently, structural studies of cilia from *Chlamydomonas*, *Tetrahymena*, and sea urchin showed that MIPs in DMT is a common feature throughout the tree of life but may have different forms in different species ^12,13^. In 2019, the 48 nm repeat structure of purified DMT from *Chlamydomonas* was solved at high resolution, and 33 types of MIPs were modeled ^14^. In 2021, the DMT structure from purified bovine trachea showed that it shares 22 conserved MIPs with *Chlamydomonas* but has a tight Tektin bundle in the A-tubule lumen that is absent in the lower eukaryotes ^15^. A further study of DMT in the human trachea revealed a similar configuration compared to bovine trachea, with 6 additional MIPs identified ^16^. These results provide an in-depth picture of DMT architecture in trachea cilia.

Unlike trachea cilia, mammalian sperm rely on vigorous ciliary beatings to overcome obstacles in the female reproductive tract. Therefore, the sperm adopt several specializations to accomplish this extreme task, such as the long axoneme, donut-shaped mitochondria in its power-generating tail, and a panoply of MIPs inside the axonemal DMT providing the stabilized force ^17-20^. It showed that overall MIP patterns in sperm and trachea differ significantly within or between species ^21^. The major differences are within the A-tubule lumen, where electron densities in the cryo-EM map of DMT in sperm axoneme are denser than trachea cilia, suggesting their different roles according to different cell motility requirements ^21^. However, the assembly details of MIPs in mammalian sperm axoneme have not yet been revealed, limiting our understanding of the structural basis of DMT in sperm beating, and the DMT-related cause of male infertility.

In this study, by using an integrative method of cryo-focused ion beam (cryo-FIB) milling, cryo-ET and STA, artificial intelligence (AI)-assisted modeling, and mass spectrometry (MS), we resolved the in-cell structure of mouse sperm axoneme DMT with a local resolution up to 3.6 Å and built the 48 nm repeat model. Then, we were able to identify 36 different mouse sperm MIPs that play the roles in reinforcing the DMT lumen, giving an insight of how mammalian MIPs weave a strong interaction network to endow the structural stability of DMT. In addition, we resolved another in-cell structure of DMT from the deformed sperm axoneme and structural comparison with the intact sperm axoneme showed a superior stability of A-tubule than B-tubule in DMT. These findings not only reveal the specialized DMT structure of mammalian sperm, but also provide a structural basis for the treatment of human infertility caused by DMT defects.

### In-cell molecular architecture of mouse sperm axoneme DMT

To elucidate the in-cell structure of DMT, we extracted mouse sperm directly from testis and vitrified them. We employed two different sample preparation approaches and collected cryo-ET datasets separately. In the first approach, sperm were vitrified on the grid with large holes (R 3.5/1) to preserve the natural state of axoneme, followed by cryo-FIB milling to prepare the cryo-lamellae with the thickness of ∼150 nm (Extended Data Fig. 1), which were used for the subsequent cryo-ET data collection (F-dataset). In the second approach, sperm were vitrified on the grid with smaller holes (R 1.2/1/3), and then the whole axoneme was used directly for cryo-ET data collection (W-dataset) at the region of sperm principal piece. In consistency with previous investigations of cilia showing multiple periodicities of DMTs ^14-16,22^, we solved three structural models of mouse sperm axonemal DMTs in the F-dataset (Extended Data Table 1) with the periodicities of 16 nm (DMT_F16_), 48 nm (DMT_F48_), and 96 nm (DMT_F96_), respectively. For the W-dataset (Table S1), we solved two structural models with the periodicities of 16 nm (DMT_W16_) and 48 nm (DMT_W48_), respectively.

In the F-dataset, the circular “9+2” architecture of the sperm axoneme can be well observed from the cross sections of selected tomograms of proper cryo-lamellae that could just hold an integral axoneme (Fig. 1A), suggesting an intact state of sperm axoneme preserved after vitrification. Although many tomograms do not contain an integral axoneme due to the limited thickness of cryo-lamellae by cryo-FIB milling ^23-25^, major components of the axoneme can be clearly observed (Fig. 1A), allowing manual picking of DMT particles in a consistent direction. To push a higher resolution, different local masks focused on A- and B-tubules were employed during image processing (Extended Data Fig. 2). The A-tubule was resolved at an overall resolution of 4.5 Å with the local resolution up to 3.6 Å in DMT_F16_, and an overall resolution of 6.5 Å in DMT_F48_ (Extended Data Fig. 3 and 4). The B-tubule was resolved at an overall resolution of 6.5 Å in DMT_F16_, and 6.6 ∼ 7.5 Å in DMT_F48_ (Extended Data Fig. 3 and 4). Based on the map of DMT_F96_, we confirmed that mouse sperm axoneme DMT has an overall periodicity of 48 nm (Extended Data Fig. 5), in consistency with the previous studies of cilia DMTs ^14,15^. For the W-dataset, similar image processing strategies were applied (Extended Data Fig. 6), resulting in maps of DMT_W16_ and DMT_W48_ with the resolutions of 7.9 Å and 8.6 Å, respectively (Extended Data Fig. 7).

**Figure 1.**
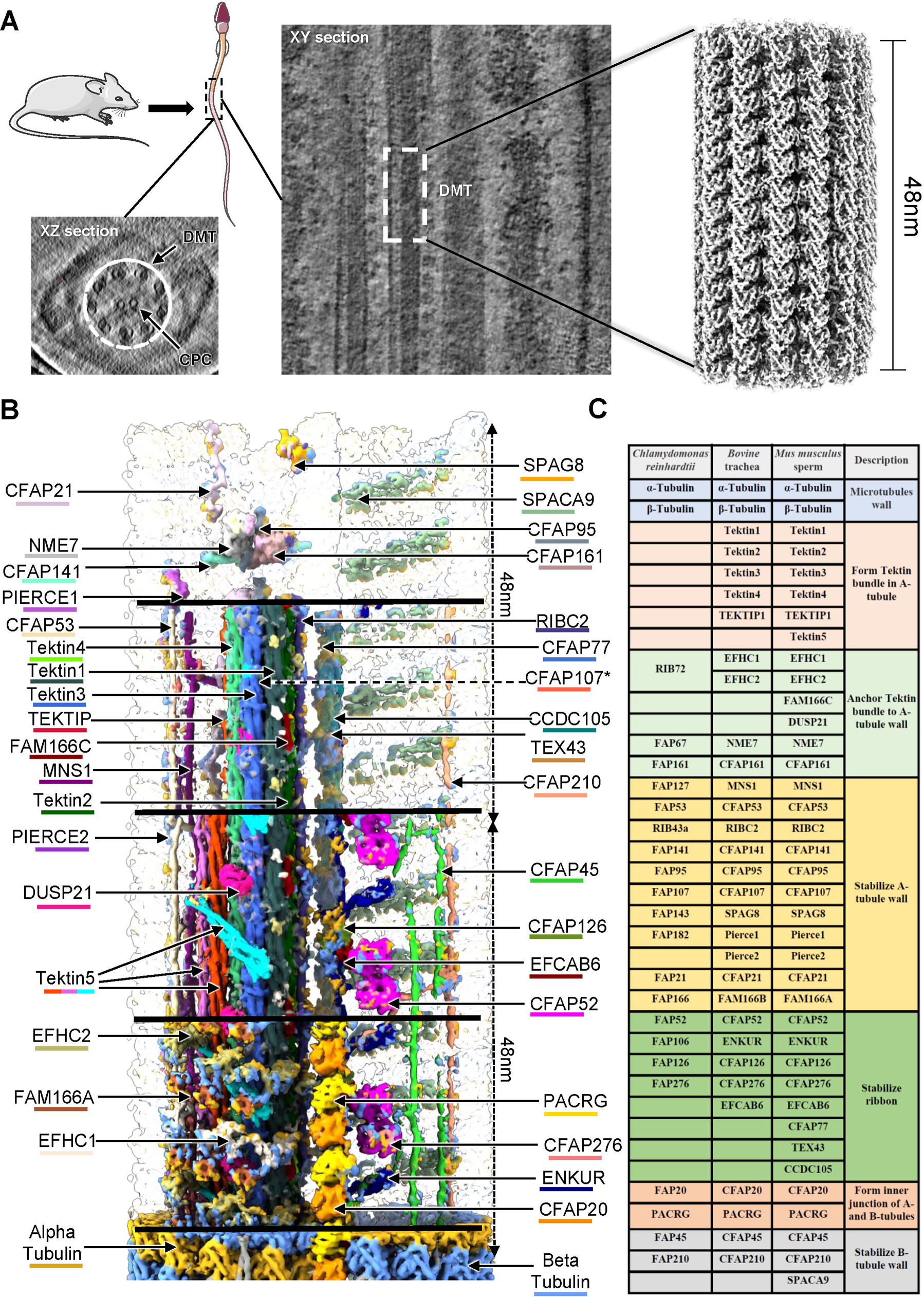
Overall architecture of mouse sperm DMT. **(A)** Schematic representation of in-cell structural determination of mouse sperm DMT. The transverse (a) and vertical (b) sections of mouse sperm axoneme are shown as the indicated slices in cryo-electron tomograms, respectively. CPC, central pair complex. DMT, microtubule doublet. A high resolution cryo-EM map of DMT with a spatial periodicity of 48 nm was obtained by sub-tomogram analysis. **(B)** Molecular dissection of DMT architecture showing with two 48 nm repeats of DMT cryo-EM map. The tubulin subunits as well as MIP subunits are colored and labeled individually. The color schemes are summarized in Extend Data Table 4. To be noted, the subunit CFAP107 is hidden by other MIPs and could not be shown in the current view. **(C)** List of names and functions of tubulins and MIPs from three representative specimen, *Chlamydomonas reinhardtii*, *Bovine* trachea, and *Mus musculus* sperm. Different colors represent different functional roles of MIPs in the architecture of DMT.

To identify multiple MIPs in the DMT map, we analyzed the proteome of mouse sperm using MS (Extended Data Fig. 8 and Extended Data Table 2). Subsequently, we employed AI-based protein structure prediction ^26,27^ and traditional manual approach to build the entire model of mouse sperm axonemal DMT from DMT_F16_ and DMT_F48_ maps (Fig. 1B and C, Fig. 2A and B). At the high-resolution region of DMT_F16_ map, well-defined side-chain densities of distinguishable residues facilitated accurate assignment and modelling of MIPs (Extended Data Fig. 9-11). For MIPs in middle-resolution area of DMT_F16_ and DMT_F48_ maps, the secondary structural elements can be resolved and the assignment and modelling of MIPs were performed with the aid by AI structural predictions and the reference by previously reported MIP structures from bovine trachea (Extended Data Fig. 12) ^15^. We used findMySequence ^27^ to identify the most possible candidate sequence of each MIP based on the complete mouse genome, yielding the successful assignments of 18 MIPs in the DMT_F16_ map with high confidence scores (Extended Data Table 3). For those MIPs not assigned by findMySequence, we started structural predictions by AlphaFold2 ^26^, refined the models based on the map and verified the model-to-map fitting quality (Extended Data Fig. 13 and 14).

**Figure 2.**
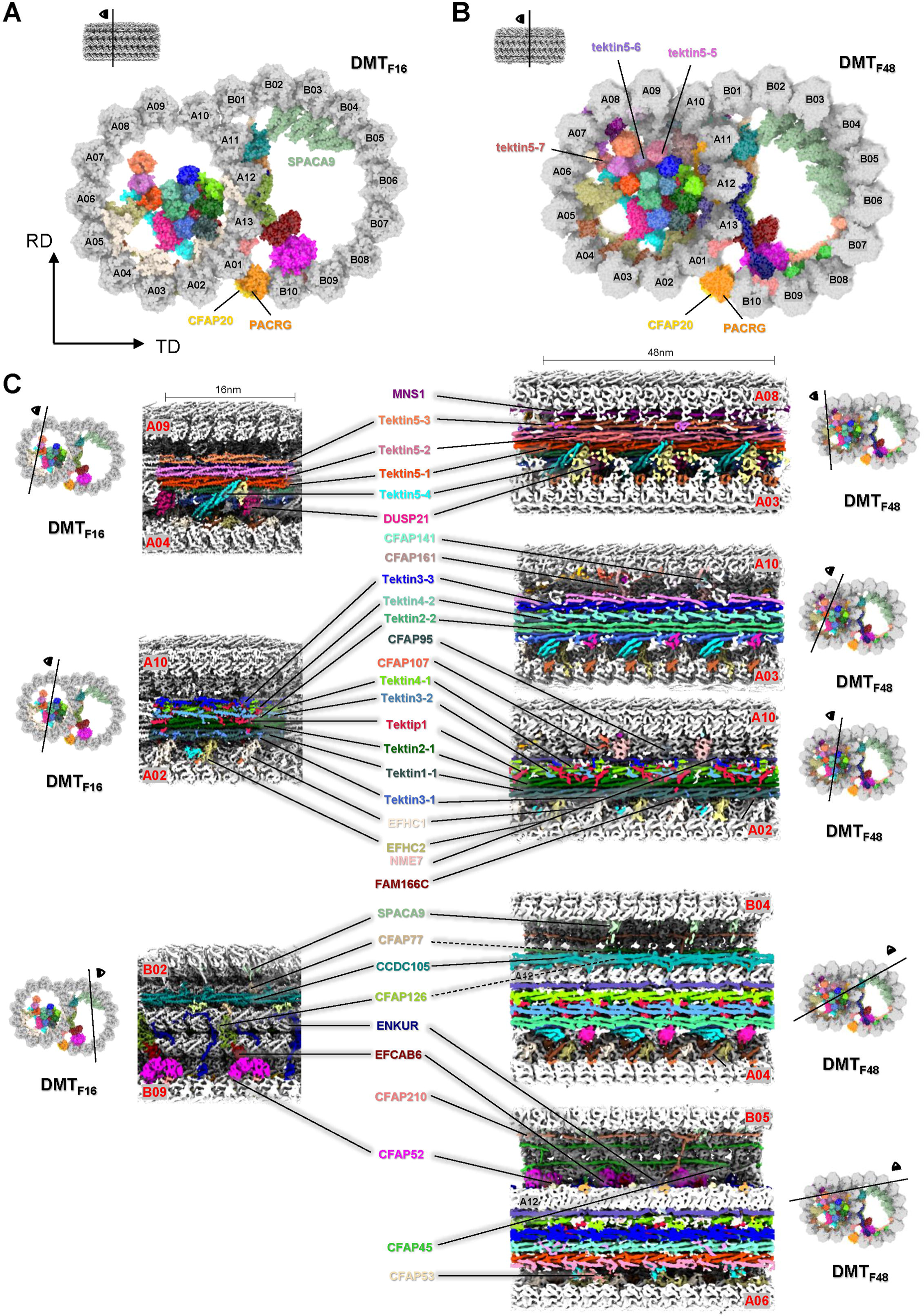
Detailed molecular organization of DMT in the repeats of 16nm and 48nm cryo-EM maps. **(A)** One representative transverse section of 16 nm repeat cryo-EM map DMT_F16_. TD, tangent direction; RD, axial direction. **(B)** One representative transverse section of 48nm repeat cryo-EM map DMT_F48_. The tubulins are numbered and MIPs are colored individually in both A- and B-tubules with CFAP20 and PACRG (A and B) and Tektin5 proteins labeled (B). **(C)** Serials of representative vertical sections of DMT cryo-EM maps in both 16nm and 48 nm repeats. The position of each vertical section is described aside. The MIPs in both A- and B-tubules are colored and labeled individually. The color schemes are summarized in Extend Data Table 4.

In addition to α- and β-tubulins, we identified and modeled 36 different kinds of mouse sperm MIPs (Fig. 1B and C, Fig. 2C). In addition to the previously reported Tektin proteins, Tektin1, Tektin2, Tektin3, Tektin4 and Tektip1 in trachea cilia, we identified and modeled a new Tektin protein Tektin5 that is specifically expressed in sperm. Besides, we identified and modeled 6 MIPs (EFHC1, EFHC2, FAM166C, DUSP21, NME7 and CFAP161) that anchor Tektin bundle to A-tubule wall, where FAM166C and DUSP21 are absent in bovine trachea cilia ^15^. The additional MIPs playing the role of stabilizing A-tubule wall include MNS1, CFAP53, RIBC2, CFAP141, CFAP95, CFAP107, SPAG8, Pierce1, Pierce2, CFAP21 and FAM166A, which are well modelled in consistency with that of bovine trachea cilia ^15^. The modelled MIPs to attach and stabilize the ribbon (defined by the protofilaments A11, A12 and A13) of DMT comprise CFAP52, ENKUR, CFAP126, CFAP276, EFCAB6, CFAP77, TEX43 and CCDC105, where CFAP77, TEX43 and CCDC105 are absent in bovine trachea cilia ^15^. The junction MIPs (CFAP20 and PACRG) between A and B-tubules as well as the MIPs (CFAP45, CFAP120 and SPACA9) attaching the inner surface of B-tubule are also well modelled, where SPACA9 was not modelled in the bovine trachea cilia ^15^.

Overall, the architecture of mouse sperm axonemal DMT is in consistency with the previous low resolution (>1 nm) study ^28^ but shows significant differences in comparison with the structure of DMT from tracheal cilia. Compared to the previously reported DMT structures from *Chlamydomonas*, *Tetrahymena*, bovine tracheal and human trachea, mouse sperm axonemal DMT structure not only exhibits an overall conserved architecture with many conserved subunits but also shows specific detailed features with many new components (Extended Data Fig. 15), indicating intrinsic structural variations between sperm axoneme and trachea cilia ^14-16,22^.

### The extended Tektin bundle in A-tubule

The Tektin family of proteins (Tektin1/2/3/4/5) play the crucial role for the formation of the axoneme (Fig. 3A). The Tektin protein possesses a triple helix bundle in its main body (Extended Data Fig. 16A), and assembles into a long filament via the canonical head-to-tail interactions (Fig. 3B). In particular, Tektin1/2/3/4 have been shown to form an 8-Tektin bundle within the A-tubule of DMT in bovine trachea cilia, contributing to the overall structural stability ^15,29-32^. Interestingly, Tektin5 is expressed exclusively in the testis and is localized throughout the sperm tail ^33^. While it was previously suggested Tektin5 is involved in the formation of the massive helix bundle inside the sperm axoneme DMT A-tubule ^34^, the exact location and copy numbers of Tektin5 were not clear due to lack of high resolution maps ^21,35^.

**Figure 3.**
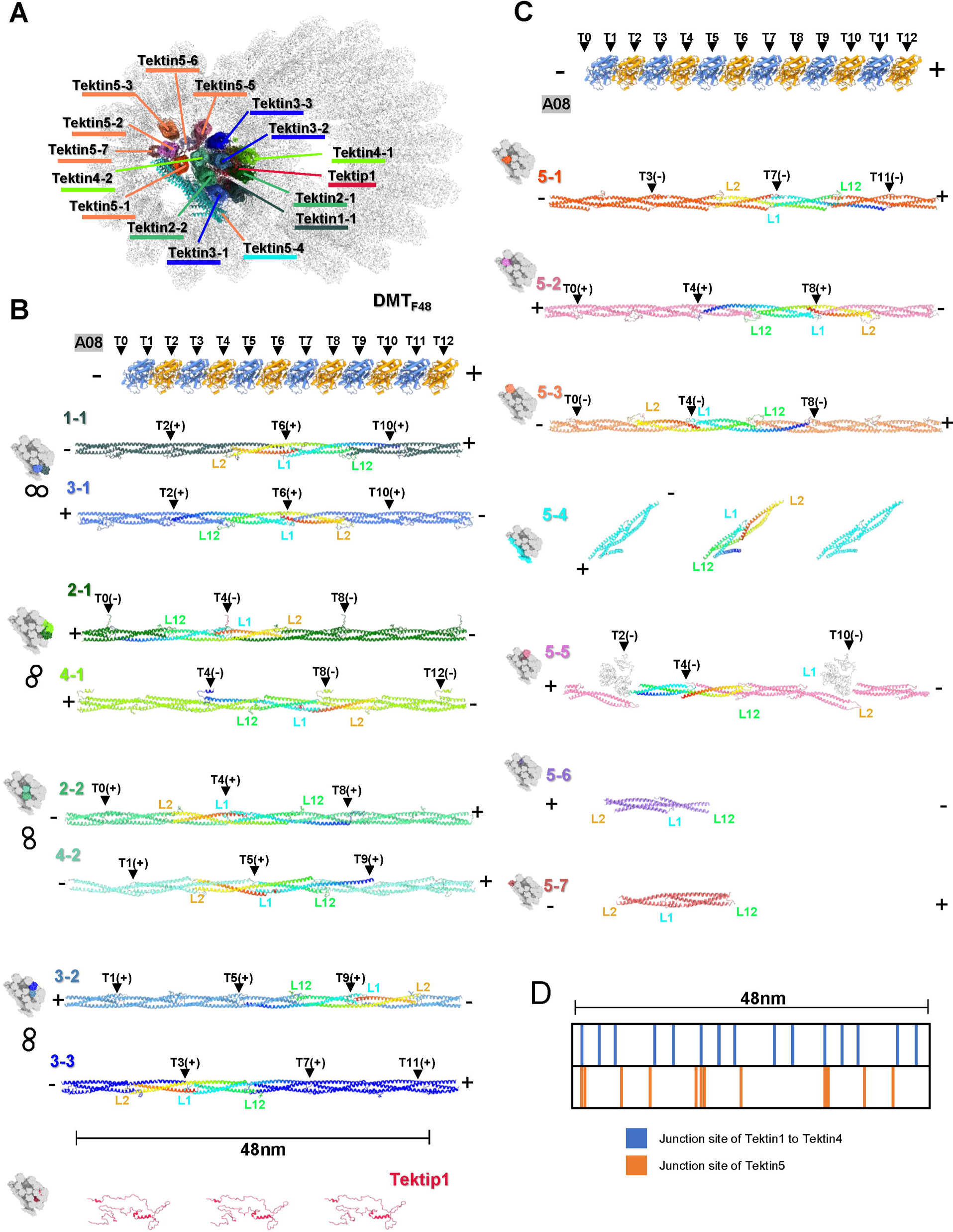
Assembly of Tektin proteins in sperm axoneme DMT. **(A) A** transverse section view of DMT_F48_ map to show the Tektin bundle in the A-tubule with Tektin proteins labelled. The coloring schemes are summarized in Extended Data Table 4. (**B**) Parallel comparisons of four pairs of Tektin1-Tektin4 filaments in each 48 nm repeat. The relative position of each pair in the Tektin1-Tektin4 bundle is shown and colored aside left. The sequential tubulins of the protofilament A08 form a 48 nm axis along the sperm axoneme with the coordinates from T0 to T12 and the direction from minus-end to plus-end of microtubule. The direction of Tektin filament is defined as the one from the C-terminus to N-terminus of one Tektin protein. The junction sites of Tektin filaments are indicated with black triangles and labelled with the coordinates Tx(+/-). Tx(+) means the site locates right to Tx and Tx(-) means left to Tx. (**C**) Parallel comparisons of Tektin5 proteins in each 48 nm repeat along with the axis defined by tubulins and with the junction sites labelled accordingly. Their positions in the Tektin bundle are also shown and colored aside left. (**D**) The distribution of junction sites of Tektin proteins in each 48 nm repeat.

Based on our high resolution in-cell DMT_16_ map with the well-resolved side chain density of many distinguishable residues, we accurately identified and modeled all the Tektin proteins inside the mouse sperm axoneme DMT. We found Tektin1/2/3/4 proteins line in a head-to-tail fashion to form homopolymer filament, four pairs of these fibers (Tektin1-1/Tektin3-1, Tektin2-1/Tektin4-1, Tektin2-2/Tektin4-2, Tektin3-2/Tektin3-3) assemble into an 8-Tektin bundle, and these fibers are organized together by the small and flexible proteins Tektip1 that form extensive interactions with the surrounding Tektin proteins (Fig. 3B), which are similar to the DMT structure in bovine trachea cilia ^15^.

Interestingly, Tektin5 seems to form an additional bundle itself (Fig. 3A and C). Domain distribution of full-length Tektin5 is basically the same as Tektin1 to Tektin4, comprising two long helices called 1A and 2A, two short helices called 1B and 2B, three loops called L1, L12, and L2 separated by these helices (Extended Data Fig. 16A) ^15^. Unlike Tektin1 to Tektin4 that share essentially the same tertiary architecture (Extended Data Fig. 16B), Tektin5 proteins adopt variant conformations (Extended Data Fig. 16C and D). Four groups of Tektin5 with the periodicity of 16 nm were identified in DMT_F16_ map, named Tektin5-1, Tektin5-2, Tektin5-3 and Tektin5-4. Among them, Tektin5-1/2/3 form a continuous homopolymer filament through the typical head-to-tail interactions (Fig. 3C). Yet Tektin5-4 adopts a completely different configuration. Instead of forming a homopolymer filament parallel to the Tektin bundle, Tektin5-4 monomer tilts about 45 degrees and lies on the outside of Tektin bundle (Fig. 3A). Each Tektin5-4 monomer is shorter than the normal Tektin protein because its N-terminal helix folds back to form a four-helix bundle (Extended Data Fig. 16C).

More interestingly, in DMT_48_ map, we identified three additional groups of Tektin5 proteins, named Tektin5-5, Tektin5-6 and Tektin5-7. There are three Tektin5-5 proteins with one in form a and two in form b (Extended Data Fig. 16C) in each 48 nm repeat (Fig. 3C). Form a and form b interact with each other in a head-to-tail way and connect indirectly with the third Tekin5-5 protein (form b) via the interactions with two NME7 proteins (Fig. 3C, see also the next section). The presence of NME7 forms a severe steric hindrance of Tektin5-5 filament, inducing a significant bending of Tektin5-5 in form b at the L2-loop (Fig. 3C). For Tektin5-6 and Tektin5-7, there is only one copy in each 48 nm repeat, respectively (Fig. 3C), locating at the opposite position of Tektin5-2 filament (Fig. 3A).

When we compared these Tektin filaments side by side along with the tubulin wall, we found their directions and junction sites are interleaved. We selected 12 sequential tubulins of the protofilament A08 to make a 48 nm axis along the sperm axoneme with the coordinates from T0 to T12, and the direction from minus-end (-) to plus-end (+) of microtubule. We further defined the direction of Tektin filament as the one from the C-terminus to (-) N-terminus (+) of Tektin protein. Then we found, for the eight-membered helix bundle core formed by Tektin1/2/3/4, half of the filaments (Tektin1-1, Tektin2-2, Tektin4-2, and Tektin3-3) have the positive direction and the another half (Tektin3-1, Tektin2-1, Tektin4-1, and Tektin3-2) with the negative direction. And their junction sites are arranged in a zigzag fashion, e.g., Tektin1-1 has the junction sites at T2, T6, and T10 while Tektin2-1 has the junction sites at T0, T4, and T8 (Fig. 3B). This specific arrangement of Tektin filaments can reinforce the tubulin wall in both directions and in multiple positions (Fig. 3D). The 7 groups of Tektin5 proteins are also arranged in a similar way. Tektin5-1, 5-3, and 5-7 orient with the positive direction, while Tektin5-2, 5-5, and 5-6 orient with the negative direction. Specially, Tektin5-4 proteins make a tilted direction. The junction sites of Tektin5 filaments are evenly distributed in each 48 nm repeat but interleaved with the junction sites of Tektin1 to Tektin4 filaments (Fig. 3D), further stabilizing the DMT architecture.

### Intricate interaction network of MIPs in A-tubule

The multiple MIP proteins in the A-tubule are bound together by a rich interaction network, which can be divided into two groups, the interactions within the Tektin bundles and the ones between the Tektin bundles and the tubulin wall. For the interactions within the Tektin bundles (Extended Data Fig. 17), the entire Tektin5 bundle anchors Tektin1/2/3/4 bundle mainly by the interactions between Tektin5-1 and Tektin4-2 (Extended Data Fig. 17A). Inside the Tektin5 bundle, we observed the interactions between Tektin5-1 and Tektin5-2 (Extended Data Fig. 17B), and between Tektin5-2 and Tektin5-3 (Extended Data Fig. 17C). Besides, inside the Tektin1/2/3/4 bundle, there are fruitful interactions between Tektip1 and Tektin2-1, 2-2, 3-1, 3-2, 3-3 and 4-1, respectively (Extended Data Fig. 17D-I), which are conserved in comparison with the interactions within Tektin1/2/3/4 bundle of bovine tracheal cilia^15^.

The interactions between the Tektin bundles and the tubulin wall are mainly mediated by multiple MIPs including EFHC1, EFHC2, FAM166C, CFAP53, CFAP141, CFAP161, NME7, SPAG8 etc., which play the roles of anchoring Tektin bundles onto the tubulin wall, providing strength along with or perpendicular to the A-tubule. (Fig. 4). It was revealed that in bovine trachea cilia EFHC1, EFHC2, RIBC2, CFAP161, NME7, and also tubulin protofilament A12 are responsible for the attachment of Tektin bundles to A-tubule wall ^15^. In our mouse sperm axoneme DMT structure, for each 16 nm repeat, we observed the interlocked interactions between the C-terminal region of Tektin5-4 and EFHC2, between Tektin1-1 and EFHC1, and between Tektin1-1 and the N-terminal region of FAM166C (Fig. 4A). To be noted, FAM166C was reported present in human trachea cilia with the role of anchoring Tektin1-1 onto the tubulin wall ^16^. For each 48 nm repeat, we observed more interlocked interactions at various transverse sections (Fig. 4B). The helical protein CFAP53 interacts with Tektin5-7 in parallel by using its second helix on one side and interacts with the tubulin protofilaments A06-A07 on the other side. Another helical protein MNS1 lies between tubulin protofilaments A07-A08 and interacts with Tektin5-3 at its C-terminal region. The junction site of two MNS1 proteins is interlocked by interacting with CFAP107 and CFAP151 that interacts with Tektin3-3. The junction sites of Tektin5-5 filaments are interlocked by the complexes of NME7/CFAP53/SPAG8 and NME7/CFAP141/CFP161, respectively. These two complexes anchor Tektin5-5 / Tektin3-3 filaments onto the tubulin wall at the sites of A07/A11 and A08/A11/A12, respectively. We also noted that the presence of NME7 with the periodicity of 48 nm hinders Tektin5-5 proteins to assemble into a long filament as Tektin5-1 to Tektin5-3, resulting in the 48 nm periodicity of Tektin5-5 filament (see also Fig. 3C).

**Figure 4.**
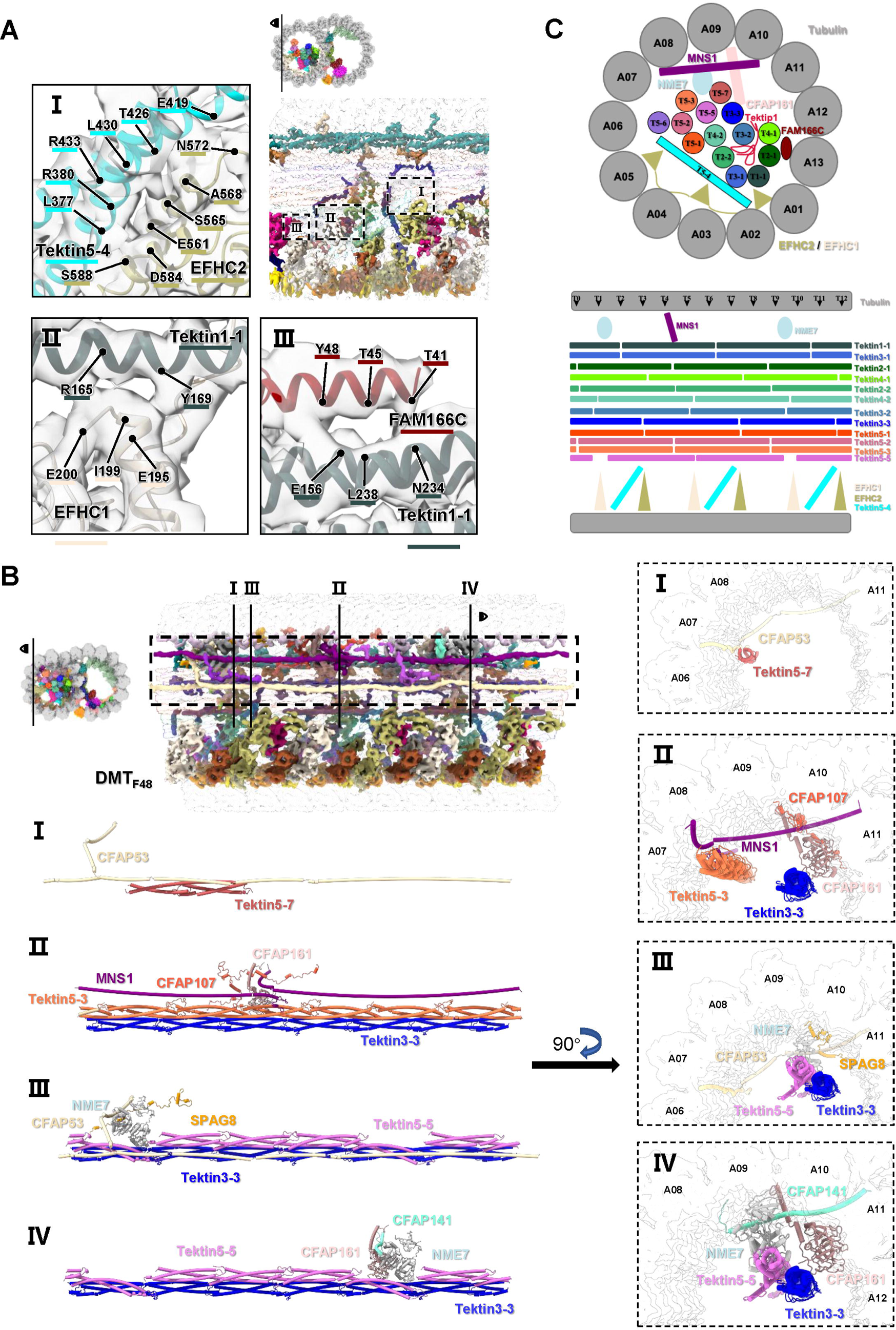
Interactions between MIPs and Tektin bundles in A-tubule and between MIPs in B-tubule. **(A)** Interactions with 16 nm periodicity. The vertical sections of DTM_F16_ map at the A-tubule and B-tubule are shown, respectively, and the MIPs responsible for anchoring Tektin bundles onto the tubulin wall in A-tubule, as well as the MIPs interacting with the ribbons in B-tubule, are highlighted and colored accordingly (Extended Data Table 4). The Tektin bundles and tubulin wall are shown in low light and transparency. The interaction interfaces with residues labelled between Tektin5-4 and EFHC2, Tektin1-1 and EFHC1, Tektin1-1 and FAM166C in A-tubule are zoomed in and shown in Panel I-III, respectively. All contacting residues were recognized using UCSF ChimeraX ^38^. **(B)** Interactions with 48 nm periodicity. The vertical sections of DTM_F48_ map at the A-tubule is shown and the MIPs responsible for anchoring Tektin bundle onto the tubulin wall are highlighted and colored accordingly (Extended Data Table 4). The Tektin bundles and tubulin wall are hidden. The interactions between Tektin5-7 and CFAP53, among Tektin5-3 filament, Tektin3-3 filament, MNS1, CFAP107, and CFAP161, among Tektin5-5 filament, Tektin3-3 filament, SPAG8, NME7, and CFAP53, and among Tektin5-5 filament, Tektin3-3 filament, CFAP141, and NME7, are zoomed in and highlighted in Panel I-IV with vertical and transverse section views, respectively. **(C)** A schematic model of MIPs interactions in A-tubule.

With the above structural analyzes, we were able to create a schematic diagram of MIPs organization within A-tubule (Fig. 4C). When viewing the junction sites of Tektin bundle, it is amazing that different Tektin filaments line up in a zigzag fashion to ensure that their junction sites are evenly distributed along the vertical axis. This kind of architecture prevents the Tektin bundle from tearing apart when the external stress is applied to A-tubule, just like a strictly weaved rope. The Tektin bundle is linked to the tubulin wall by several buffer proteins, like EFHC1/EFHC2, MNS1 and NME7 with different periodicities, to reinforce the instability of tubulin wall. From the view of transverse section (Fig. 4C), the tilted Tektin5-5 provides additional supporting force for the Tektin bundle, EFHC1 and EFHC2 reinforce the protofilaments A01 to A05, FAM166C for the protofilament A13, and MNS1 for the protofilaments A08 to A10. As a result, MIPs in A-tubule are efficiently assembled and work together to reinforce the structural stability of DMT.

### Organization and interaction of MIPs in B-tubule

The B-tubule is formed by the protofilaments B01 to B10, which surround the ribbon region defined as the protofilaments A11 to A13 ((Fig. 2A and B) ^36^, and the gap between the protofilaments A01 and B10 is bridged by the inner junction formed by PACRG and CFAP20 (Fig. 1B and 2A). PACRG and CFAP20 are mostly formed solely by α helices and β sheets, respectively (Fig. 5A). Both proteins are highly conserved throughout eukaryotes, from *Chlamydomonas* to humans, and have the same function in mediating interaction of A- and B-tubule ^14-16,37^.

**Figure 5.**
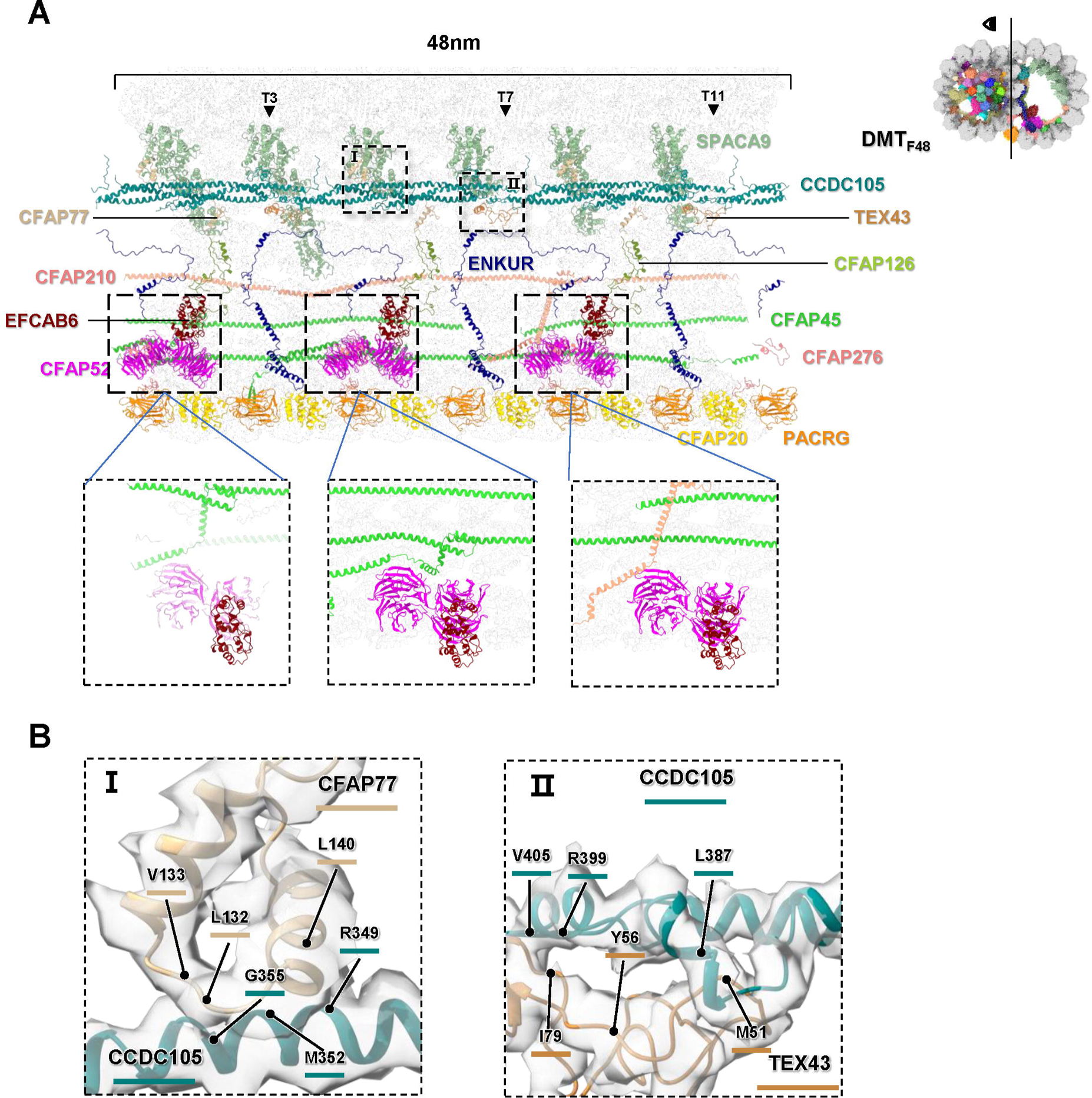
Structure organization of MIPs in B-tubule. **(A)** Structural models of all MIPs in the B-tubule are shown with a 48 nm length of vertical section nearby the ribbon region, labelled and colored accordingly (Extended Data Table 4). The coordinates (T3, T7 and T11) of junction sites of CCDC105 filament are indicated. The interactions between CFAP52/EFCAB6 in 16 nm repeat and CFAP45/CFAP276 in 48 nm repeat are zoomed in. **(B)** The interaction interfaces between CCDC106 and CFAP77, and CCDC105 and TEC43 in B-tubule, which are indicated in (A), are zoomed in and shown in Panel I and II with residues labelled, respectively.

Compared to the highly decorated A-tubule, B-tubule possess less MIPs, including CCDC105, TEX43, CFAP77, CFAP126, CFAP276, EFCAB6, CFAP52 and ENKUR (Fig. 5A), which attach to the ribbon and extend into the B-tubule lumen. Besides, there are additional MIPs including SPACA9, CFAP45 and CFAP210, which decorate the inner wall (B02-B10) of B-tubule (Fig. 5A and Extended Data Fig. 9).

As one of the largest proteins attached to the ribbon, CCDC105 comprises a three-helix bundle in its main body with an additional helix at its N-terminus (Extended Data Fig. 16E). Structural superposition reveals CCDC105 resembles a

similar domain configuration with Tektin proteins and the r.m.s.d. of C-alpha atoms between CCDC105 and Tektin4-1 is 1.35 Å that is calculated by UCSF ChimeraX ^38^. CCDC105 resides between the protofilaments A11 and A12, assembling into a long head-to-tail filament by interacting with the neighbor CCDC105 molecules. The coordinates of junction sites of CCDC105 filament are T3, T7 and T11, respectively (Fig. 5A). At the junction site, we observed there is a small protein TEX43 clamping the L12 loop (L387-R399) of CCDC105 (Fig. 5A and B). It also interacts with the tubulin wall at the A12 protofilament, acting as a rivet to strength the junction site of CCDC105 filament (Fig. 5A). Another small protein CFAP77 also interacts with CCDC105 at the region of M352-G355 to stabilize the ribbon (Fig. 5B).

ENKUR that has an extremely elongated configuration (Fig. 5A) connects the protofilaments A12, A13, and B10 in every 16 nm repeat, and participates in the inter-protofilament interactions of B-tubule. We observed a potential interaction between the middle region of ENKUR and the loop region of TEX43, suggesting a role of reinforcement.

EFCAB6, the MIP with no ortholog in *Chlamydomonas*, was reported to attach to the protofilament A13 in the B-tubule of bovine trachea cilia DMT with 48 nm periodicity ^15^. Interestingly, in our structure of mouse sperm axonemal DMT, we found EFCAB6 with the periodicity of 16 nm, each EFCAB6 interacts with CFAP52, which contains two WD40 domains and attaches to the CFAP45 or CFAP210 filaments (Fig. 5A). The additional copies of EFCAB6/CFAP52 pairs provide a stronger support to the protofilaments A13 and B10, in comparison with that of trachea cilia ^15,39^ (Fig. 5A).

Both CFAP45 and CFAP210 proteins exhibit a 48 nm periodicity and possess long helices to form filaments attaching to the protofilaments B07-B09 (Fig. 2B, C and 5A) ^39^. The two copies of CFAP45, crawling on protofilaments B08 and B09 in parallel, adopt different tertiary configurations in the N-terminus. The N-terminal helix in one CFAP45 contains an elbow bend to accommodate the steric hindrance of CFAP52 (Fig. 5A), and the same helix in another CFAP45 protein are plain helices. CFAP210 also contains an elbow bend at its N-terminal helix across protofilaments B07 to B09 to accommodate the steric hindrance of CFAP52 (Fig. 5A), while its rest part lies between protofilaments B06 and B07.

The most outstanding features of B-tubule are the pyramid-shaped densities decorated on its inner surface at the region of protofilaments B02 to B06 (Fig. 2B). Both recent studies on human sperm microtubule singlets (SMT) and trachea DMT showed that it is SPACA9 with a pyramid-shaped structure to decorate both lumens of DMT B-tubule and SMT with the same repeat pattern ^16,40^. In the DMT of human trachea cilia, SPACA9 was reported to have an 8 nm periodicity and situated in between the α- and β-tubulin heterodimers ^16^. At every 8 nm, three SPACA9 monomers are situated between the protofilaments B02 to B05, forming a hierarchy structure ^16^. However, in the DMT structure of mouse sperm axoneme, we observed that SPACA9 forms a pattern of 5-3-3-4-4-4 in each 48 nm repeat, and SPACA9s are located between the protofilaments B02 to B07, B02 to B05, and B02 to B06, respectively (Fig. 5A and Extended Data Fig. 9). Since SPACA9 is located at the cross-sectional position between tubulin heterodimers and different protofilaments, these abundant SPACA9 proteins provide additional stabilization to the B-tubule walls. To be noted, although SAXO1 was found to mediate the interaction between SPACA9 and the tubulin wall in DMT of human trachea cilia ^41^, we did not identify the equivalent in our structure, which might be due to the limited low resolution of the local region.

### DMT structure in a deformed sperm axoneme

Since there are more MIPs in A-tubule than that in B-tubule (Extended Data Fig. 18), A-tubule would exhibit a better structural stability than B-tubule. The cryo-ET dataset (W-dataset) using the sperm specimen vitrified on the grids with a smaller hole (R1.2/1/3) resulted the tomograms with the deformed sperm axoneme (Fig. 6A), providing us an opportunity to investigate the role of MIPs in maintaining the structural stability of sperm axoneme DMT. In this dataset, the “9+2” architecture of the sperm axoneme is compressed along the electron illumination direction, possibly due to the surface tension applied to the sperm tail during blotting. By analyzing the cryo-EM maps of DMT_W48_ and DTM_F48_ (Fig. 6A and B), in addition to the different widths, 432 Å for DMT_W48_ and 443 Å for DTM_F48_, we further found that the A-tubule in the deform specimen (DMT_W48_) shows a similar structure as that in the intact specimen (DTM_F48_), while the densities of B-tubule in the deformed one (DMT_W48_) are significantly averaged out (Fig. 6C and D).

**Figure 6.**
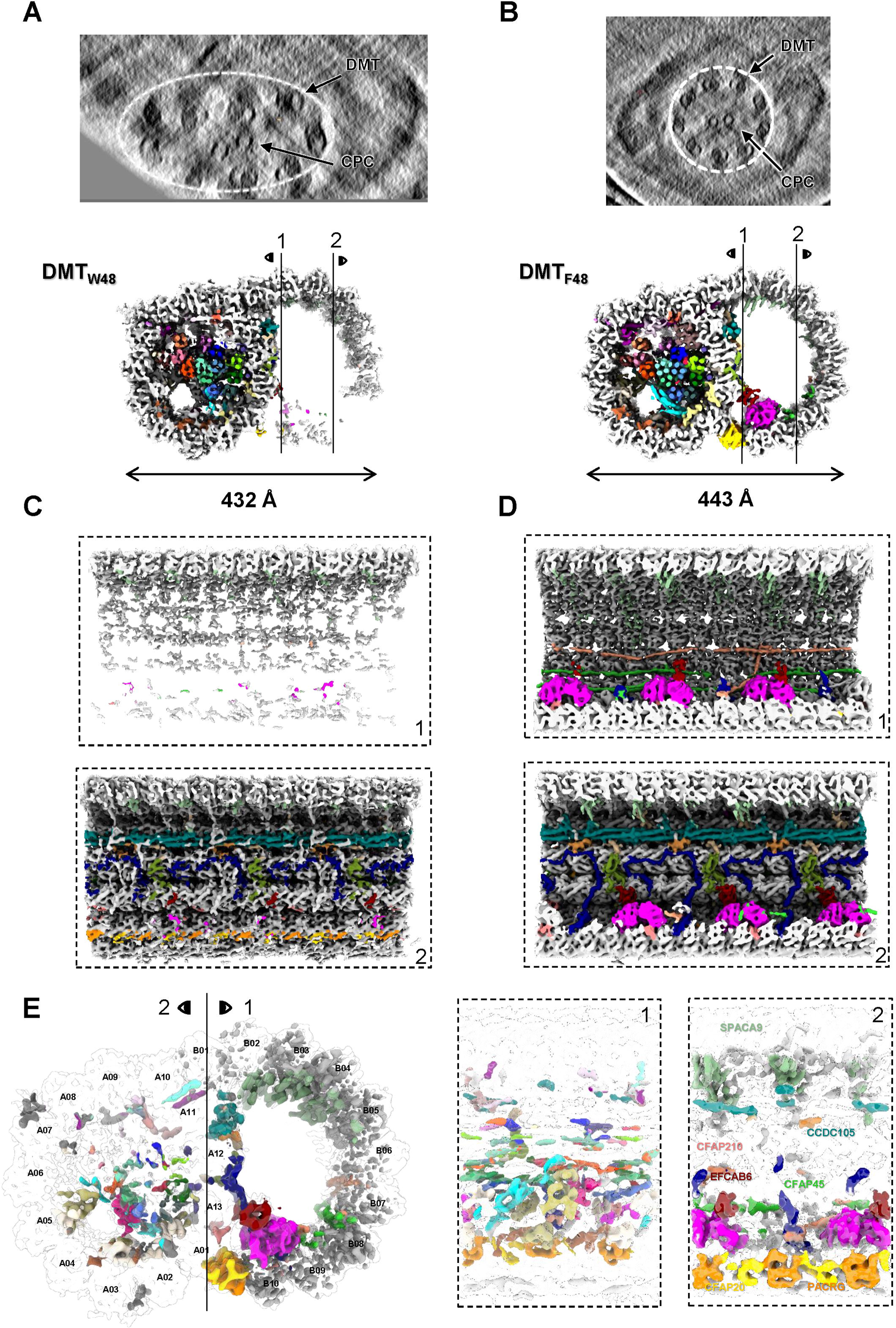
Comparison of DMT structures between deformed (DMT_W48_ map) and intact (DMT_F48_ map) axonemes of mouse sperm. **(A) and (B)** Selected tomograms of W-dataset (A) and F-dataset (B) are shown in XZ sections with DMT and CPC labeled. The transverse sections of DMT_W48_ (A) and DMT_F48_ (B) maps are shown with MIPs densities colored, respectively. **(C) and (D)** Two vertical sections of DMT_W48_ and DMT_F48_ maps are shown with MIPs densities colored, respectively. The section positions (1 and 2) are indicated in (A) and (B), accordingly. **(E)** Difference map by subtracting DMT_W48_ map from DMT_F48_ map with the transverse and vertical sectional views. The densities corresponding to different MIPs were colored accordingly (Extended Data Table 4).

To compare the structural changes of DMT between the deformed and intact states, we calculated the difference map by subtracting DMT_W48_ from DTM_F48_ (Fig. 6E). From the difference map, we observed small chunk of densities of the tubulin wall of A-tubule, suggesting the 13 protofilaments of A-tubule exhibit the same structure in both datasets. In the A-tubule lumen, although the Tektin bundle between two specimens remains unaffected with debris of densities, we observed significant difference densities in the bridging components such as EFHC1/2 and DUSP 21 that are responsible for anchoring the Tektin bundle to the tubulin wall (Fig. 6E). It suggests that the Tektin bundle provides the main role to support the A-tubule stability when responding the external forces, and the bridging components may disperse or buffer the external forces through appropriate conformational changes to minimize the deformation of the tubulin wall.

However, in the B-tubule, large amounts of tubulin walls and MIPs are present in the difference map, including CFAP20、PACRG、SPACA9 and CFAP52 (Fig. 6C, D and E), indicating that these proteins are the most fragile part of sperm axonemal DMT whose structures are the most vulnerable to external forces. The inner junction formed by CFAP20 and PACRG is probably the most stressed position in the deformed sperm DMT, because the density of inner junction has almost disappeared in DMT_W48_ map (Fig. 6A). On the other side of the B-tubule, the densities of tubulin walls were preserved (Fig. 6C and D), which may be due to the interaction between A10 and B01 protofilament and the reinforcement bySPACA9 proteins.

As a result, by comparing the sperm axonemal DMT structures in the intact and deformed states, we found MIPs in DMT play different roles in maintaining the overall structures of A- and B-tubules. The different compositions of MIPs result in a great difference in the stability of A- and B-tubules. The super stability of A-tubule is due to the abundance of Tektin proteins that form Tektin bundle in the lumen.

## Discussion

In this study, we solved the in-cell molecular structure of mouse sperm axonemal DMT with a local resolution up to 3.6 Å and built 36 different types of MIPs within the 48 nm repeat. We found, in mouse sperm axonemal DMT, the lumen of A-tubule is almost fully occupied by various MIPs, while the B-tubule is relatively hollow. A series of Tektin5 proteins in different conformations reinforce the helix bundle composed by Tektin1 to Tektin4, and other MIPs with different periodicities form intricate interaction networks to anchor the Tektin bundle onto the tubulin wall of A-tubule. In B-tubule, we found the MIPs like CCDC105/TEX43/CFAP77 attaching to the ribbon and SPACA9 bound to the inner wall play the roles to reinforce the structural stability of B-tubule. More importantly, by comparing DMT structures from the intact and deformed mouse sperm axonemes, we revealed the importance of MIPs to resist external force, where A-tubule due to its intricate interaction networks of MIPs exhibits a significant stronger structural stability than B-tubule. The super stability of A-tubule is necessary to serve as a structural base for ODA, IDA, RS and N-DRC binding, which is highly essential to ensure that sperm can complete the complex swimming mode in the reproductive tract. These results advance our knowledge of mammalian sperm structure as well as our understanding of the roles of MIPs in sperm axoneme and shed the light on the future investigation of disease associated mutations of MIPs.

Compared to the trachea cilia, sperm has an extremely long axoneme for swimming in the reproductive tract. In addition, the length of sperm flagella varies across different species, such as ∼ 60 μm for Sea urchin, ∼ 60 μm for human and ∼ 120 μm for mouse ^23,42^. The difference in the length of flagella would be related to fit with the specific environment during the fertilization process and longer flagella would result in a better swimming power and better ability of fertilization. With the longest axoneme, mouse sperm requires the DMT of its axoneme a hyper structural stability. Our in-cell structural study provides such molecular insight into the hyper structural stability of mouse sperm axonemal DMT and reveals rich MIPs as well as intricate interaction networks in the A-tubule lumen, which are quite different from sea urchin sperm ^43^. Thus, the long-term evolutionary selection would lead to a direct positive correlation between the sperm axonemal length and the abundance of MIPs within DMT. In line with this, the axoneme of sperm is longer than that of trachea cilia, and the viscosity of the reproductive tract mucus is higher than that of the respiratory tract mucus, suggesting that sperm faces greater resistance during swimming ^20,44^. Therefore, it needs a more stable MIPs assembly system to protect the entire axoneme from external force and complete the fertilization process.

During the preparation of this manuscript, we noticed a preprint study on the isolated DMT of bovine sperm using cryo-EM single particle analysis (cryo-SPA) ^45^. Interestingly, we observed that the MIPs identified in mouse sperm DMT were also present in bovine sperm DMT, indicating a conserved axonemal structure in mammalian sperm. However, in bovine sperm, the SPACA9 exhibited a 48 nm periodicity with five groups of four SPACA9 proteins situated between the protofilaments B02 to B06, and one group of five SPACA9 proteins located between the protofilaments B02 to B07, forming a 5-4-4-4-4-4 pattern ^45^, which is in contrast to the 5-3-3-4-4-4 pattern observed in mouse sperm DMT here, suggesting a species-specific arrangement of SPACA9, which is worthy of further investigation. Moreover, the mouse sperm DMT contains fewer Tektin5 proteins than the bovine counterpart. In mouse sperm DMT, only one Tektin5-6 protein is present in each 48 nm repeat, while two copied are found in bovine sperm (referred to as Tektin5E) ^45^, warranting further studies.

In our DMT_F96_ map, densities attributable to the IDA, ODA, N-DRC, RS1, RS2, and RS3 can be identified, enabling further understanding of the complete structure and motion mechanism of the sperm axoneme. With the further efforts including technology developments, it is foreseeable that a complete high resolution “9+2” architecture of the sperm axoneme will be resolved, providing in-depth understanding into molecular mechanism of sperm swimming and genetic diseases of infertility.

## Methods

### Mouse sperm extraction

To extract mouse sperm, 200 μl sperm capacitive solution (Extended Data Table 5) was taken as drops in a 35 mm dish, covered with mineral oil, and placed at 37 °C in a 5% CO_2_ incubator for more than half an hour. 12-week-old C57 male mice were sacrificed for cervical dislocation, and the epididymis was dissected. The sperm was taken out under a stereoscope and the sperm pellets were placed in a capacitive solution. The sperm pellets were placed in the incubator for more than half an hour until the sperm pellets were completely dispersed. The sperm solution was taken out and placed in 1.5 ml Eppendorf tube and temporarily stored on ice before further experiments.

The animal experiments were performed in the Laboratory of Animal Center of Institute of Biophysics, Chinese Academy of Sciences, in accordance with the National Institutes of Health Guide for the Care and Use of Laboratory Animals and according to guidelines approved by the Institutional Animal Care and Use Committee at Institute of Biophysics with Dr. Guangxia Gao as chairman of the board.

### Sample preparation for the intact axoneme of mouse sperm

Freshly extracted sperms were centrifuged under 4 °C, 400 G (Thermo Scientific Legend Micro 17R) for 5 min. The sediment of every 100 μl of the sperm solution was re-suspended carefully into 100 μl pre-cooled PBS on ice, then diluted 5.5 times with PBS right before use. Cryo-EM grid (Quantifoil R3.5/1, Au 200 mesh) was glow discharged using Gatan Solarus for 60 s. The sperm sample was vitrified using Leica EM GP or Leica EM GP2. 2.7 μl of PBS diluted sample was applied onto the grid, followed by immediate blotting for 2∼5 s at 100% relative humidity and 4 °C, then plunge frozen into liquid ethane cooled to −186 °C, then stored in liquid nitrogen before cryo-FIB milling.

Sperm lamellae were prepared using a Aquilos 2 SEM (ThermoFisher Scientific). First, the sample was coated with a platinum layer for 10 s, followed by a coat of organometallic platinum layer for 15 s, then followed by a final coat of platinum layer for 10 s using default settings. After selecting optimum positions for milling, a stepwise tilting and milling manner was performed to obtain final lamellae: (1) tilt to 10°, set current to 300 pA, mill a 1.5 μm gap of 13 μm in width, (2) tilt to 10.5°, set current to 100 pA, mill a gap of 800 nm from the topside and of 12.75 μm in width, (3) tilt to 9.5°, mill a gap of 500 nm from the downside and of 12.75 μm in width, (4) tilt to 10°, set current to 50 pA, mill a gap of 300 nm from both side and of 12.25 μm in width, (5) set current to 30 pA, and with extensive tilting to 10.2° or 9.8° to reach a final gap of 80 nm from both side with 12 μm in width. All tilt angle aforementioned are defined as the relative tilting angle with respect to the grid. Once reached the optimum thickness, the grid was sputtered with a final layer of platinum at 30 KV, 10 mA for 1 to 3 s, then stored in liquid nitrogen before data collection in TEM.

### Sample preparation for the deformed axoneme of mouse sperm

Freshly extracted sperms were centrifuged under 4 °C, 400 G (Thermo Scientific Legend Micro 17R) for 5 min. The sediment of every 200 μl of the sperm solution was re-suspended carefully into 100 μl pre-cooled PBS on ice, then diluted 20 to 40 times with PBS before further experiments. Cryo-EM grid (Quantifoil R1.2/1.3, Au 200 mesh) was glow discharged using Gatan Solarus for 60 s right before use. The diluted sperm solution was mixed at a ratio of 1:1with 10 X concentrated 10 nm protein A-coated gold fiducials (Electron Microscopy Sciences). 2.7 μL of the mixture was applied onto the grids and blotted for 1.5 to 2.5 s in 100% relative humidity and 4 °C, then plunged into liquid ethane for vitrification, and transferred into liquid nitrogen before data collection, this process was performed by using Vitrobot Mark IV (Thermo Fisher Scientific).

### Cryo-ET tilt series acquisition

Both cryo-FIB milled and directly frozen grids were mounted onto Autoloader in Titan Krios G3 (Thermofisher Scientific) 300 KV TEM, equipped with a Gatan K2 direct electron detector (DED) and a BioQuantum energy filter. Tilt series were collected under a magnification of 81,000 X, resulted in a physical pixel size of 1.76 Å, or 0.88 Å under super-resolution mode, in K2 DED. Before data collection, the pre-tilt of the sample was determined visually, and the pre-tilt was set to be 10° or −9° to match the pre-determined geometry caused by loading grids. The total dose was set to 3.5 electrons per square angstrom per tilt, fractioned to 10 frames in a 1.2 s exposure, and the tilt range were set to be between −66° to +51° for −9° pre-tilt or −50 to +67° for +10° pre-tilt, with a 3° step, resulting in 40 tilts and 140 electrons per tilt series. The slit width was set to be 20 eV, with the refinement of zero-loss peak after collection of each tilt series, and nominal defocus was set to be −1.8 to −2.5 μm. For directly frozen sperm samples, the tilt range was set to be −60° to +60°, starting at 0°, with all other parameters unchanged. All tilt series used in this study were collected using a dose-symmetry strategy-based beam-image-shift facilitated acquisition scheme, by in-house developed scripts within SerialEM software ^46-48^.

### Image processing of F-dataset

After data collection, all fractioned movies were imported into Warp for essential processing including motion correction, Fourier binning by a factor of 2 of the super-resolution frames, CTF estimation and tilt series generation ^49^. Then the tilt series were subjected to AreTOMO for automatic tilt series alignment ^49,50^. The aligned tilt series were inspected visually in IMOD, and misaligned frames were discarded to generate new tilt series in Warp. This process was repeated three rounds in total, and the tilt series with less than 30 frames or completely failed in alignment were kept from further processing ^50,51^. The alignment parameters of all remaining tilt series were transferred back to Warp, and initial tomogram reconstruction was done at a pixel size of 10 Å in Warp.

DMT particles was manually picked using a filament picking tool in Dynamo, by picking the start and end points of each DMT filament and separating each crop point by 8 nm along the filament axis ^52^. The 3D coordinates and two of three Euler angles (except for in-plane rotation) were generated by Dynamo, then transferred back to Warp for exporting of sub-tomograms ^49,52,53^. Sub-tomogram refinement was done in RELION 3.0 or 3.1 ^54,55^. First, all particles were reconstructed at a pixel value of 14.08 Å, and an initial reference was generated by low-pass filtering at 60 Å of directly average of all picked-out particles, then a K=1 3D classification with restrictions of the first two Euler angles was done for 100 iterations, and particles within 16 nm were discarded using Dynamo to avoid overlapping. After alignment on the 14.08 Å level, the aligned parameters were transferred back to Warp to export sub-tomograms at a pixel size of 7.04 Å, then one round of 3D auto refinement was done in RELION, with a mask covering solely the A-tubule of 16 nm repeat of DMT.

From this point on, the data processing was split into different branches. For 16 nm repeat maps, the aligned parameter was transferred back to M for multi-particle refinement, and the resolution reaches the Nyquist limit. Then the sub-tomograms corresponding to A- and B-tubule were reconstructed at 3.52 Å pixel size by M, and extensive refine and 3D classification were performed in RELION ^56^. When resolution of these particles stopped increasing, the aligned parameters were transferred back to M to export A- and B-tubule sub-tomograms at binned 1 level, then extensive auto refinement in RELION and M refinement was done, which reached a final resolution of 4.5 Å for A-tubule MIP and 6.5 Å for B-tubule.

For 48 nm repeat maps, a mask inside A-tubule lumen of 16 nm repeat map was applied find the map with 48 nm periodicity, then sub-tomograms of 3.52 Å pixel size were exported for auto refinement in RELION. Then, six different sets of masks were applied evenly onto A- and B-tubule, and auto refinement in RELION and M refinement was done using these masks, reaching final resolutions of 6.5 to 7.5 Å. 96 nm repeat maps were generated by classifying 48 nm repeat map with a local mask covering N-DRC region (Extended Data Fig. 2-5).

### Image processing of W-dataset

The data processing of W-dataset is basically the same as that of F-dataset. The 16 nm repeat map was generated by auto refinement at binned 2 level in RELION and M refinement to reach a final resolution of 7.9 Å. The 48 nm repeat map was auto-refined in RELION and further refined in M with a final resolution of 8.6 Å (Extended Data Fig. 6-7). The data processing statistics were summarized in Extended Data Table 1.

Mask creation at different image processing stages was done by a combination of UCSF Chimera, Dynamo, and RELION, and converting of Dynamo table file or RELION star file or tilt series alignment files was done by a wrap of automatic tools of sub-tomogram data processing ^52-55,57,58^.

Visualization of cryo-EM maps, image making, and rendering of refined models was performed using either UCSF Chimera, UCSF ChimeraX, or 3D Protein Imaging^38,58,59^.

For difference map calculation, DiffMap in CCP-EM software package were used^60^.

### Mass spectrometry

Mouse sperm for mass spectrometry identification was cleaned with PBS and then centrifuged at 2000g for 5 min. We then treated the sperm with five different buffers (B1: 0.1% Triton; B2: 0.6M NaCl; B3: 8M Urea; B4: 10% SDS; B5: 0.1% Triton+8M Urea). For 10% SDS, we added the buffer to sperm precipitation and heated it at 95 ° C for 5 min. The other buffers were incubated at room temperature for 10 min after sperm precipitation. The treated sperm was centrifuged at 2000g for 10 min, and 20µl supernatant was taken for SDS-PAGE. Coomassie brilliant blue was stained and decolorized using eStain L1 protein stain instrument (L00657C) (Extended Data Fig. 8).

After decolorization, the strips were enzymolized with trypsin overnight, and then the peptides were extracted with acetonitrile of different concentrations in multiple steps. The peptide mixture obtained by enzymolysis was analyzed by liquid chromatography-tandem mass spectrometry. Then SEQUEST HT search engine of Thermo Proteome Discoverer (2.4.1.15) was used to search and identify proteins in the Uniprot_proteome_mouse (update-07/2022) protein library database. The search results were authenticated by mass spectrometry on nanoLC-Orbitrap Exploris 480.

### Model building

Modelling was based on each asymmetric unit of tubulin dimer and MIPs. Since there were no available structures of mouse DMT proteins in the PDB database, the AlphaFold ^61^ predicted models were initially used as the starting model. Three different strategies were used to generate the final model.

First, for proteins in DMT_F16_ map with considerably higher local resolution and map quality, initial poly-alanine models were manually built into the density map using Coot ^62^, then findMySequence program ^27^ was employed to search for the protein candidate within the mouse protein database (17,141 reviewed sequences in UniProtKB). Upon identification of a high-confidence protein, the predicted AlphaFold structure was fitted into the density. The model was then manually adjusted using COOT and further refined via pheix.real_space_refine in Phenix ^63^. This process was iterated several times to generate the final model. The models of tubulin α/β, CCDC105, CFAP20, CFAP276, CFAP52, DUSP21, EFCAB6, EFHC1, EFHC2, PACRG, Tektin1 to Tektin4, Tektin5-1, Tektin5-2, and TEKTIP1 were all built in this manner (Extended Data Fig. 11 and Table 3). For α-tubulin family proteins, which have several outputs in the findMySequence program, we chose the highest-ranked protein candidate for modeling based on the MS results list (Extended Data Dataset 1).

Second, for proteins exhibiting lower resolution in local regions, the findMySequence program failed to identify meaningful candidate. We referred to the reported DMT structure in bovine trachea ^15^ to assign the homologous protein, and flexibly fitted the predicted AlphaFold structure to the density. Subsequently, we manually adjusted the model using COOT and further refined it using pheix.real_space_refine in Phenix ^63^. This process was iterated several times to generate the final model. The models of NME7, CFAP161, CFAP161, MNS1, CFAP53, RIBC2, CFAP141, CFAP95, CFAP107, SPAG8, Pierce1, Pierce2, CFAP21, FAM166A, ENKUR, CFAP126, CFAP45, and CFAP210, were all built using this approach.

Lastly, for proteins with lower resolution and not involved in the bovine trachea DMT structure, we selected the top 1500 candidates from the MS results list (Extended Data Dataset 1). We manually fitted their AlphaFold-predicted models into the local density to identify the best match. For densities with multiple matching candidates, we selected the highest-ranked protein based on the MS results list. Subsequently, we adjusted the model using COOT and refined it with pheix.real_space_refine in Phenix ^63^. This process was iterated several times to generate the final model. The models of remaining Tektin5 proteins (5-3, 5-4, 5-5, 5-6, 5-7), FAM166C, CFAP77, TEX43, and SPACA9, were all built using this approach.

### Data availability

The sub-tomogram averaged cryo-EM maps for A tubule in 16 nm repeat, B tubule in 16 nm, A tubule (part1, 2, 3) in 48 nm repeat, B tubule (part1, 2, 3) in 48 nm repeat, section I of DMT in 96 nm repeat, section II of DMT in 96 nm repeat as well as the composite maps DMT_F16_, DMT_F48_ and DMT_F96_, in F-dataset have been deposited in the Electron Microscopy Database (EMDB) under the accession codes EMD-35210, EMD-35211, EMD-35222, EMD-35224, EMD-35225, EMD-35226, EMD-35227, EMD-35228, EMD-35231, EMD-35232, EMD-35229, EMD-35230 and EMD-35236, respectively. The sub-tomogram averaged cryo-EM maps for DMT_W16_ and DMT_W48_ have been deposited in EMDB under the accession codes EMD-35237 and EMD-35238, respectively. The atomic models of DMT in 16 nm repeat and in 48 nm repeat have been deposited in Protein Data Bank with the accession codes of 8I7O and 8I7R, respectively.

## Acknowledgments

We thank Ping Shan, Lianwan Chen, and Tongming Zhang (F.S. laboratory) for their assistance in laboratory management. We would like to thank Center for Biological Imaging (CBI), Institute of Biophysics (IBP), Chinese Academy of Science (CAS) for the support of specimen vitrification, cryo-FIB milling, and cryo-ET data collection, and we are grateful to Dr. Shuoguo Li, Lulu Qin, and Dr. Jianguo Zhang for their help with sample preparation and data collection. We are particularly grateful to Prof. Yao Cong (Shanghai Institute of Biochemistry and Cell Biology, CAS) for her valuable advice on the project, and Dr. Alister Burt (MRC-LMB, UK) for his help on image processing. We would like to thank Dr. Dongdong Fan (IBP, CAS) and Dr. Jifeng Wang (IBP, CAS) for their help in sperm preparation and mass spectrometry experiment, respectively.

This work was equally supported by grants from National Natural Science Foundation of China (31925026 to FS), National Key Research and Development Program (2021YFA1301500 to FS), the Strategic Priority Research Program of the Chinese Academy of Sciences (XDB 37040102 to FS). This work was also supported by grants from National Natural Science Foundation of China (32071187 to YZ and 31830020, 61932018 to FS), and National Key Research and Development Program (2019YFA0904101 to YZ).

## Author contributions

Y.Z., and F.S. conceived and coordinated the study; L.T., G.Y., Y.Z., and F.S. designed the research; L.T., G.Y., and X.H. performed the experiments; L.T., G.Y., and Y.Z. analyzed the data; L.T., and G.Y. wrote the manuscript; Y.Z., and F.S. revised the manuscript.

